# Preventions of oral self-mutilation in patients with Lesch-Nyhan syndrome

**DOI:** 10.1101/425967

**Authors:** Lorevanda V. de Carvalho, Luan F. de Oliveira, Tiago F. Chaves

## Abstract

Lesch-Nyhan syndrome (LNS) is a rare genetic disorder linked to the X chromosome that implies an inborn error of metabolism, due to deficiency in the purine metabolic pathway, affected individuals present neurological symptoms, such as cognitive and motor impairment, besides behavioral changes such as self and hetero-aggression. This characteristic aggressive behavior leads to insults and self-mutilation, thus requiring methods of self-protection, some invasive and permanent, such as tooth extraction. Noninvasive methods, such as the use of oral protectors, pharmacological therapies and botulinum toxin A applications, are also employed to decrease discomfort and the implications of such behavior. We describe briefly the methods of prevention against oral self-mutilation in patients with Lesch-Nyhan syndrome found in the literature.

## INTRODUCTION

First described in 1964, Lesch-Nyhan syndrome (LNS) is a rare disorder linked to the X chromosome, that approximately one in 380,000 live births (Nyhan (1997)). It causes disturbance in the uric acid metabolism and central nervous system (CNS)(Lesch and Nyhan (1964)) due to mutations in the gene that codes for the hypoxanthine-guanine phosphoribosyltransferase enzyme (HPRT1), located between Xq26 and Xq27, an important enzyme in the purine rescue route (Bell et al. (2016)). This results in total or partial lack of HPRT activities and high level of uric acid concentration in urine (Dabrowski and Medicine (2005)). The pathology may also be associated with dopaminergic activity deficits in the basal ganglia (Visser et al. (2000)). The characteristic framework of LNS consists of a set of neurological behaviors and symptoms, presenting a delay in the motor development and variable degrees of cognitive impairment, self-injurious behavior (self-mutilation) that manifest during the first year of life (Bell et al. (2016)). Patients are generally not able to sit independently and remain impassable throughout life, more specifically, self-mutilation, which generally leads to the correct diagnosis of Lesch-Nyhan syndrome (Christie et al. (1982); Schroeder et al. (2001)). According to Anderson and Ernst (1994), out of the 40 Lesch-Nyhan patients, the study found that 90% had permanent physical damage due to self-mutilation. Patients are aware of their self-harming behavior and become agitated and afraid when their protective and restrictive gear are removed (Lesch and Nyhan (1964)). The restriction method seems to be the most suitable for controlling self-mutilation. Regarding the restrictions and nozzle protections used to avoid self-mutilation, tooth extraction has been the most used method to reduce self-harming behavior (Campolo González et al. (2018)). Even after tooth extraction, these edentulous patients turn to other forms of self-mutilation, in many cases using their fingers to cause self-injury (Gutierrez et al. (2008)). Other noninvasive methods include mild mouthguard use and psychiatric pharmacological therapy to avoid further damage to soft perioral tissues. Recent treatments include the use of botulinum toxin type A (BTX-A) injected into the bilateral masseters. BTX-A temporarily prevents presynaptic release of acetylcholine, causing motor plaque dysfunction and muscle weakness (Dabrowski and Medicine (2005); Zilli and Hasselmo (2008); Jeong et al. (2006); Olson and Houlihan (2000)). The focus of this work is to present prevention methods against oral self-mutilation in patients with Lesch-Nyhan syndrome found in literature.

## PREVENTIVE METHODS

### Pharmacotherapy

Among the treatments used to alleviate the causes of this syndrome is the use of drugs aimed at inhibiting the conversion of guanine and hypoxanthine to uric acid by HPRT. One of the most commonly used drugs is allopurinol, treatment with allopurinol reduces serum levels of urate and uric acid and thus prevents crystaluria, nephrolithiasis and gout. Some benzodiazepines and gamma-aminobutyric acid inhibitors are used for the treatment of spasticity and dystonias, but due to the lack of knowledge about neurological dysfunctions, useful therapies have not been found, and rehabilitation and muscle and postural training programs are recommended. And for the control of self-mutilation, neuroleptics are used to minimize this behavior. Benzodiazepines and carbamazepine are the most useful for decreasing anxiety and suppressing harmful behavior (Zilli and Hasselmo (2008); Jeong et al. (2006)). Despite the numerous benefits always reported to the use of these medications (Campolo González et al. (2018); Gutierrez et al. (2008)). Buitelaar (1993), advises caution for many reasons. Primarily due to the sedative properties induced by these drugs that prevent patients in both their cognitive functions and their physical abilities, and secondly due long-term use side effects. In a case reported by Jeong et al. (2006), the patient was receiving a daily dose of 2 mg diazepam to relieve the self-injurious action, but their success was limited. After being submitted to a new psychiatric consultant, his medication was changed to sertraline 12.5 mg and 0.25 mg risperidone, to control anxiety and self-injurious behavior. After seven days the dosages of both drugs were doubled. After 15 days, the self-mutilation and shaking behavior was significantly reduced and the interaction between the patient and his mother improved. According to the author within 1 month, the frequency of self-injurious behavior was limited to less than a single attempt per week, after 4 months of such pharmacotherapy resulted in the complete disappearance of self-injurious behavior without any side effects.

### Oral Device

The use of mouthguards is widely described in the technical literature. The use of mouth guards is extensively described in the technical literature were described by Anderson and Ernst (1994); Jeong et al. (2006). Scott and Ranalli (2005), mentions that the applications of oral protection devices are many and varied, from the use as protector to the therapeutic one. Olson and Houlihan (2000), describe the follow-up of a case, which the pediatric dentist build a two-piece acrylic mouth guard to fit firmly on the patient’s upper and lower teeth. In addition, there was a verbally reinforcing training for the patient to keep the mouth guard in the correct mouth position. Jeong et al. (2006), emphasizes in his case study the development of a soft mouthguard, which was fabricated and modeled using Biostar (Scheu-Dental, Iserlohn, Germany). The upper dental arch was covered with a mouth guard made of soft resin material. During the periodic dental monitoring phase, the patient appeared to be well adapted to this soft mouthguard, and reduced frequency of general self-mutilation included fewer lip bites in the first 2 weeks of mouthguard use along with pharmacotherapy, according to the author’s emphasis the treatment results were very promising. The most commonly used oral device is a soft mouth guard (Dabrowski and Medicine (2005); Jeong et al. (2006); Olson and Houlihan (2000)).

1. Bite-Blocks (Figure 1);
2. A combination of extraoral and intraoral orthodontic apparatus that covers the chin and is held in place with a rubber band around the head or with a helmet on the neck-strap (Figure 2);
3. Acrylic trays intended to force the lower lip anteriorly onto the lower teeth (Figure 3B);
4. Various types of shields that protect the tongue and lips from direct injury (Figure 3A);
5. Labial bumpers welded onto orthodontic bands or stainless-steel crowns (Figure 4);

**Figure 1.**
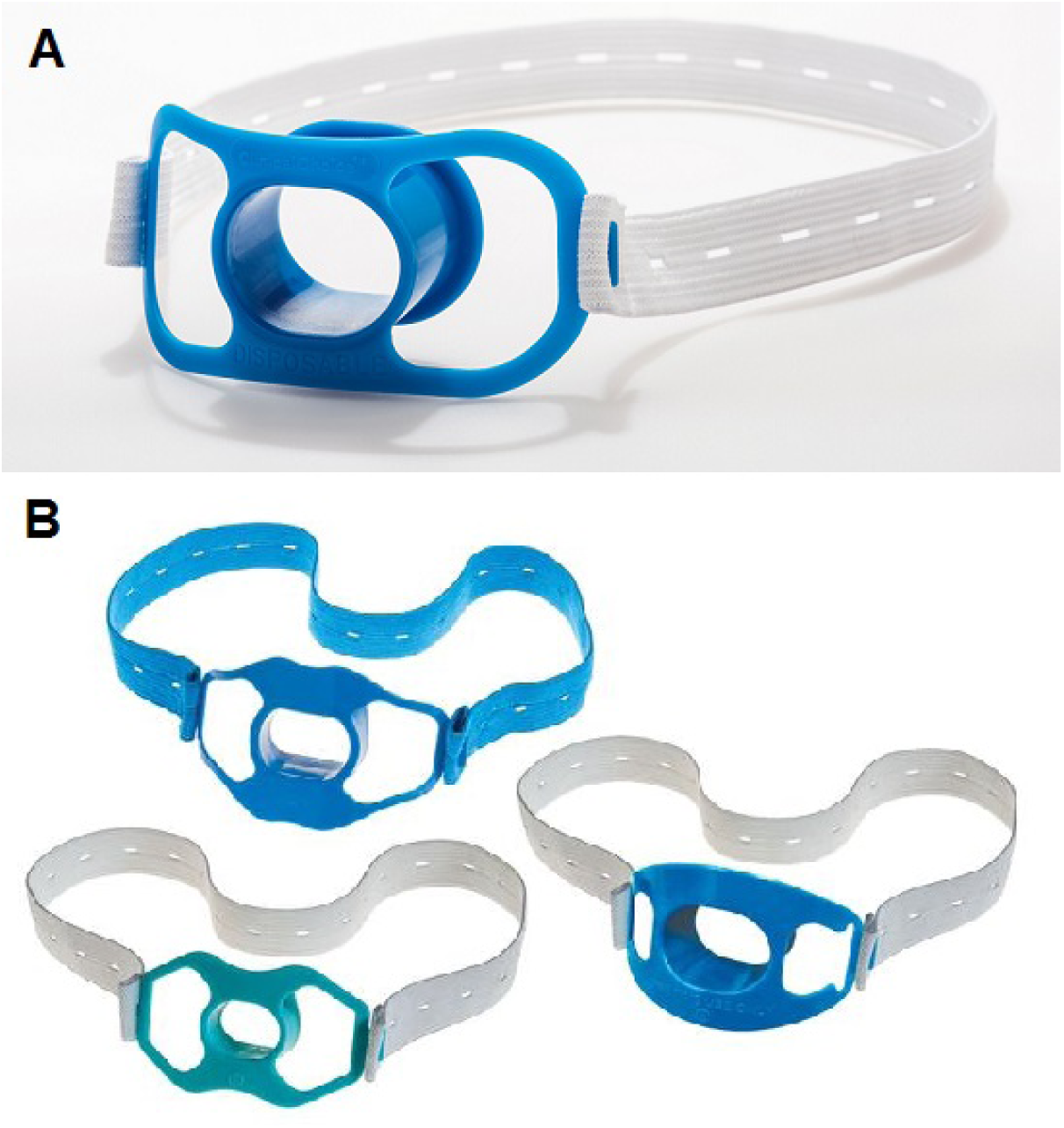
Models of Bite Blocks. (A) Endoscopic Bite Blocks by Choice, (B) ScopeValet™ Endoscopic Bite Blocks by Healthcare (2018).

**Figure 2.**
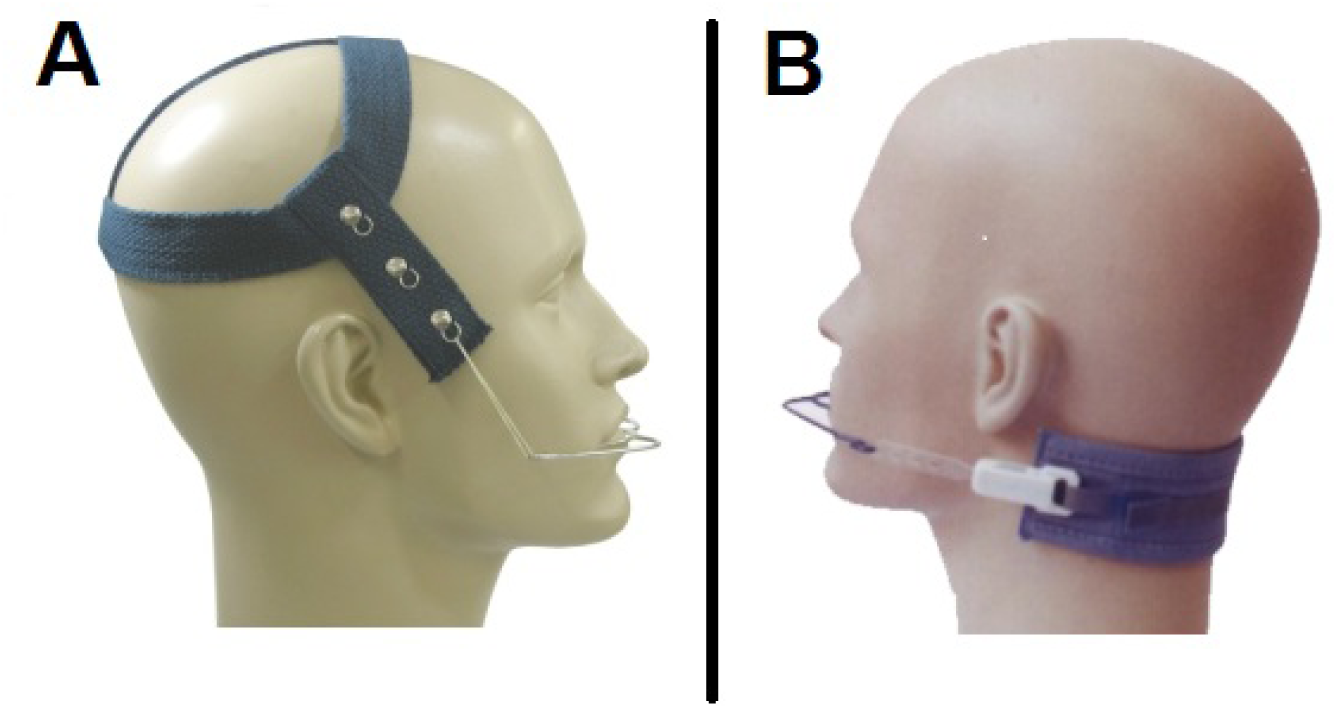
Models of helmet on the neck-strap. (A) Extra Oral Supplies by Biodental (2018), (B) Extraoral Supplies by JJORTHO and Tencent (2018).

**Figure 3.**
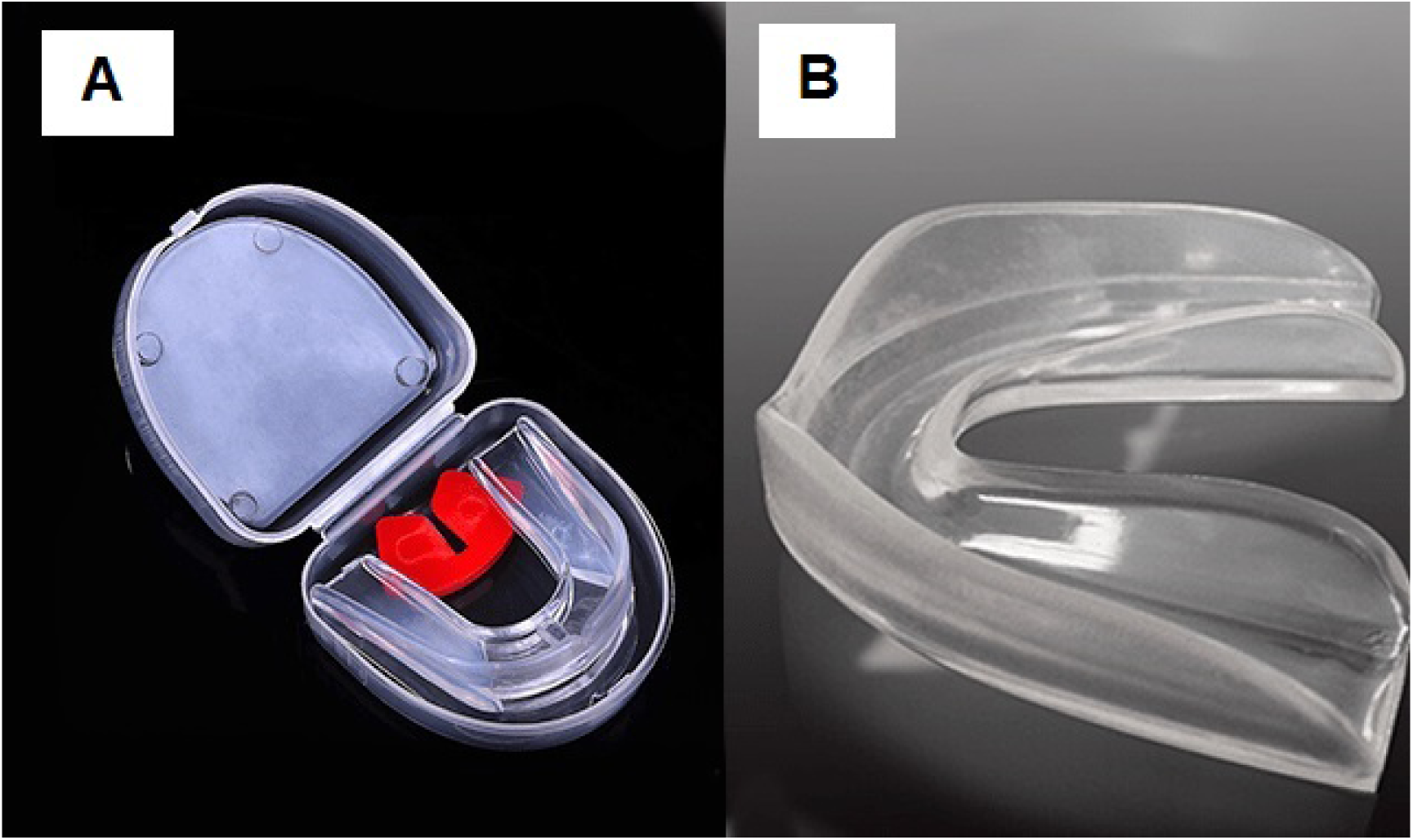
Mouthguard double model (A) that is used at the bottom and top and mouthpiece singular model (B) with single format for upper teeth. Source: Fighter (2018).

**Figure 4.**
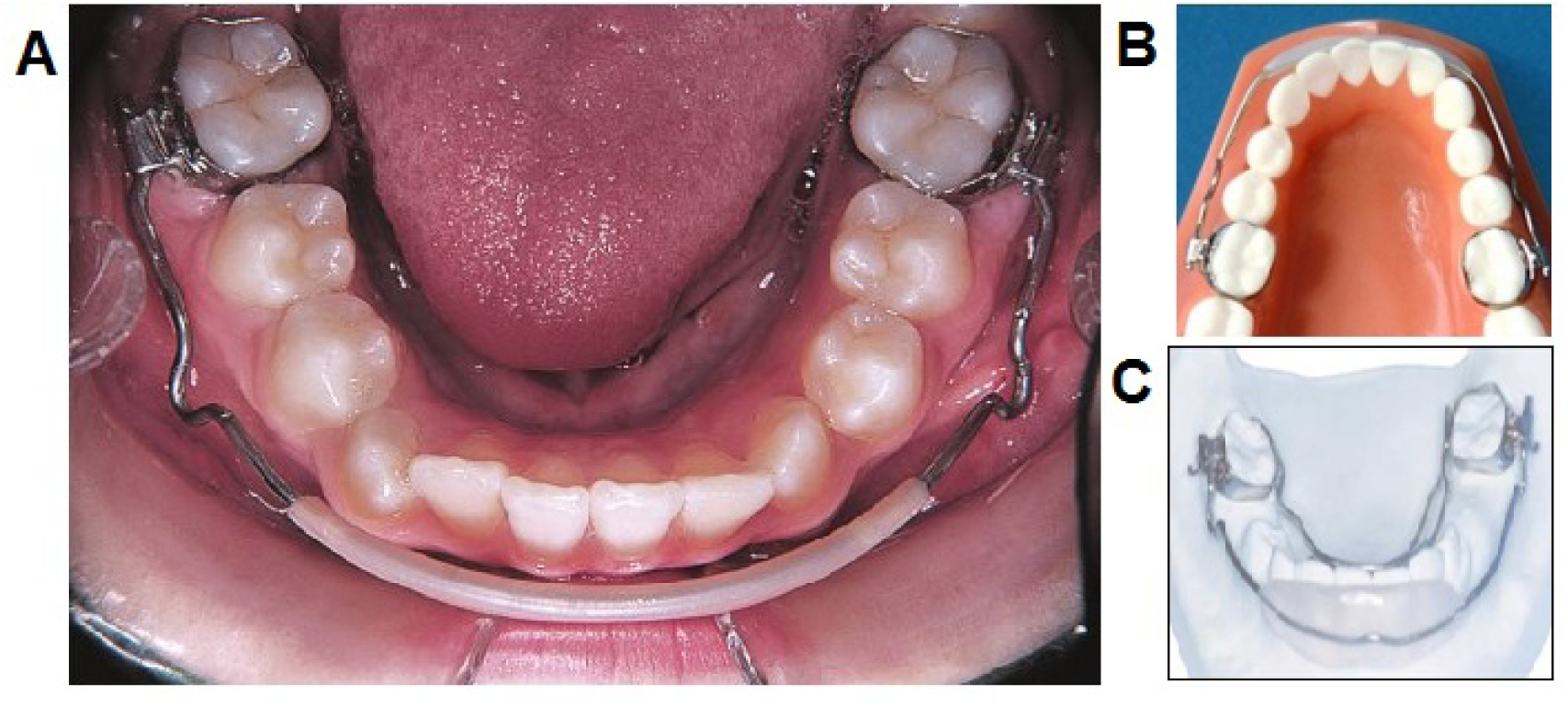
(A) Intraoral occlusal view with hard arc labial bumpers (Jacob et al. (2014)). (B and C) Lip Bumpers Models (Care (2018)).

### Botulinum Toxin Type-A

Dabrowski and Medicine (2005), present a case of an LNS patient who underwent botulinum toxin A (BTX-A) treatment injected into the bilateral masseters. The active principle of BTX-A is to temporarily prevent presynaptic release of acetylcholine, causing motor plaque dysfunction and muscle weakness. As reported, the treatment resulted in a significant reduction in self-injurious behavior and healing of the patient’s local lesions. This work suggested a biological mechanism of action of BTX-A in the central and peripheral nervous system, which results in a reversal of this behavioral pathway. In the treatment of the mentioned study, BTX-A was administered in both masseter muscles, with 20 units of BTX-A being injected at each site, meaning a total of 40 units per muscle group. The toxin was reconstituted with 1cm3 of saline solution; its application was performed using a 27-gauge needle and 1cm3 syringe. These applications were repeated every 3 months for a series of three visits. BTX-A was well tolerated, and benefits lasted up to 10 weeks, when at last the behavior returned, requiring the use of restrictions. During this treatment the patient was able to return to school, his wounds on his hands, lips and tongue healed completely, his speech ability improved, although he continued to be abnormal. According to the authors, no impact was observed on the ability to eat or swallow, which contributed to the patient’s physiology. The family emphasized a decrease in self-harm behavior as well as hostility. The conclusion of the treatment suggested that BTX-A may be more active, either directly on the peripheral nervous system or indirectly on the central nervous system, interfering with the pathway of behavior that causes self-mutilation to occur in the LNS. Other study carried out by Gutierrez et al. (2008), reports the application of BTX-A on the muscles of a 30-year-old male patient. According to Guttierrez, the application of BTX-A in muscles instead of masseters is due to the patient having his lesions mainly on the lips. The author of the study also points out that the other method employed by Dabrowski and Medicine (2005) requires significant doses of BTX-A and may have spread to the pharyngeal muscles, thus affecting dysphagia. In this methodology 2 sessions were applied every 3 months, the injections in the both zygomatic muscles, in the lower part of the orbicularis oculi muscle and lower upper eyelid of the lip. During applications, there was a significant drop in oral self-mutilations. This study reinforces the results of the therapeutic approach of BTX-A, reporting a decrease in harmful behavior, improvement of patient’s tissues and no adverse side effects. Despite the limitations of these studies, because it is applied in only two invidious, it is a safe and effective alternative therapeutic treatment for the decrease or elimination of the self-mutilations in LNS(Dabrowski and Medicine (2005); Gutierrez et al. (2008)).

## FINAL CONSIDERATIONS

The technical literature concludes that there are no specific references methods for the prevention of this type self-mutilation, methods need to be developed and applied according to the needs of each patient. As an alternative to dental extraction, it is suggested a combination psychiatric therapy, pharmacological, dental treatment, personalized mouthguard and, in extreme cases, the application of treatment based on botulinum toxin type-A is a viable option in reducing, decreasing and in eliminating self-injurious behavior involving the oral pathways.

## AUTHOR CONTRIBUTIONS

All authors contributed equally to the elaboration, revision and approval of the final article.

